# Investigating Fibroblast-Induced Collagen Gel Contraction Using a Dynamic Microscale Platform

**DOI:** 10.1101/628230

**Authors:** Tianzi Zhang, John H. Day, Xiaojing Su, Arturo G. Guadarrama, Nathan K. Sandbo, Stephane Esnault, Loren C. Denlinger, Erwin Berthier, Ashleigh B. Theberge

## Abstract

Mechanical forces have long been recognized as fundamental drivers in biological processes, such as embryogenesis, tissue formation and disease regulation. The collagen gel contraction (CGC) assay has served as a classic tool in the field of mechanobiology to study cell-induced contraction of extracellular matrix (ECM), which plays an important role in inflammation and wound healing. In a conventional CGC assay, cell-laden collagen is loaded into a cell culture vessel (typically a well plate) and forms a disk-shaped gel adhering to the bottom of the vessel. The decrement in diameter or surface area of the gel is used as a parameter to quantify the degree of cell contractility. In this study, we developed a microscale CGC assay with an engineered well plate insert that uses surface tension forces to load and manipulate small volumes (14 µL) of cell-laden collagen. The system is easily operated with two pipetting steps and the microscale device moves dynamically as a result of cellular forces. We used a straightforward one-dimensional measurement as the gel contraction readout. We adapted a conventional lung fibroblast CGC assay to demonstrate the functionality of the device, observing significantly more gel contraction when human lung fibroblasts were cultured in serum-containing media versus serum-free media (p≤0.01). We further cocultured eosinophils and fibroblasts in the system, two important cellular components that lead to fibrosis in asthma, and observed that soluble factors from eosinophils significantly increase fibroblast-mediated gel contraction (p≤0.01). Our microscale CGC device provides a new method for studying downstream ECM effects of intercellular cross talk using 7-35 fold less cell-laden gel than traditional CGC assays.

## Introduction

Fibroblasts are key mesenchymal cells in connective tissue which synthesize extracellular matrix (ECM) components and provide structural support for the extracellular environment (Kendall and Feghali-Bostwick, 2014). As part of the tissue self-repair mechanism, fibroblasts interact with surrounding ECM proteins through a variety of inflammatory mediators and differentiate into a more contractile phenotype known as myofibroblasts (Jeffery, 2001; Royce et al., 2012). However, overreactive myofibroblasts generate and deposit excessive ECM proteins in the interstitium, contributing to fibrotic diseases such as asthma and idiopathic pulmonary fibrosis (Grinell, 2003; Hinz et al., 2007). Therefore, understanding the mechanobiology of fibroblasts in ECM and the underlying signaling mechanisms is essential to developing therapies for diseases involving fibrosis. The goal for this study is to develop a microscale assay that captures and reflects dynamic fibroblast-ECM interactions.

The fibroblast-induced collagen gel contraction (CGC) assay was established by Bell et al. to study fibroblast-matrix interactions (Bell et al., 1979) and has been modified and widely used over the past four decades. The traditional CGC assay is performed by embedding fibroblasts into a three-dimensional (3D) gel matrix, such as collagen or fibrin, on the bottom of a well plate, which is then manually separated from the well plate surface (for example by scraping a pipette tip around the perimeter of the well) to loosen the gel puck from the well plate walls and enable contraction (Dallon and Ehrlich, 2008; Mikami et al., 2016). The contractile forces generated by fibroblasts propagate throughout the collagen matrix and arrange collagen fibers to higher density structure with decreased matrix volume (Jonas and Duschl, 2010). As a result, measuring the decrease in size of a gel matrix puck by imaging and subsequent analysis provides a direct way to assess fibroblast contractility.

Addressing some complications in the existing CGC assay workflow could help researchers meet a diverse set of experimental needs. For example, deformation of gel shape and ambiguous post-contraction gel borders make the exact gel area difficult to define (Chen et al., 2013); the requirement of relatively large cell samples precludes the assay from use with limited primary cells (Redden and Doolin, 2003); large volumetric consumption of gels (>100 μL per replete in a 96 well plate) is relatively expensive (Gullberg et al., 1990; Timpson et al., 2011); and the friction between gel and substrate upon gel contraction is not well defined, potentially adding variation to the data (Chen et al. 2008; Vernon and Gooden, 2002). Through the years, numerous tools and technologies have been developed to improve these shortcomings. For example, automated image analysis programs have been used to improve accuracies for the geometric gel shape readout (Jin et al., 2015; Chen et al., 2012). Leung et al. developed a high-throughput microscale aqueous two-phase droplet fabrication method in conventional 384-well plate which effectively reduced the gel droplet to 10-15 µL (Leung et al., 2015). Ilagan et al. used glass capillary to cast cell-laden collagen which was subsequently detached from the glass surface by pipetting force, significantly reducing friction and converting three-dimensional parameters into a single, linear measurement (Ilagan et al., 2009). The recent efforts to develop new CGC assay platforms have underscored the utility of the assay and motivated our work to develop a new microscale CGC assay that builds on past improvements and enables a combination of new experimental features.

Our goal was to create a microscale CGC assay that addresses the needs in objective gel shape quantification, reducing cell and gel consumption, minimizing friction between the plastic culture substrate and the gel, and enabling segregated coculture to study paracrine signaling. Here, we describe a microscale CGC platform based on 24 well plate insert that enables the study of dynamic cell-matrix interactions. The device is characterized by a two-step pipetting operation and a simple angle measurement as a contractility readout. We demonstrate a proof of concept use of this technology with a serum stimulation experiment. Further, we validate our device for coculture by testing the hypothesis that soluble factors secreted by eosinophils induce increased gel contraction by lung fibroblasts as has been previously observed using the traditional CGC assay (Zagai et al., 2004). In the future, we envision that we and other researchers could use our technology to address additional research questions relating to paracrine signaling in lung fibrosis as well as fibrosis in other organs such as the regulation of epithelial-to-mesenchymal transition in kidney fibrosis and the role of stellate cell activation in hepatic fibrosis.

## Materials and Methods

### Device fabrication

Devices were fabricated using a Form 2 SLA 3D printer (Formlabs, Somerville, MA). 3D-printed devices were designed with Solidworks and converted to .form files with PreForm 2.11.0 (Formlabs) prior to being printed with Form 2 Clear V4 Resin (Formlabs). After printing, devices were sonicated in isopropanol (IPA) for 15 min, rinsed with fresh IPA, and UV-cured (Quans 20 W UV Lamp) for 2 hours. Original design files are included in the ESI.

### Cell culture

Human fetal lung fibroblasts (HFL-1) were obtained from the American Type Culture Collection (Rockville, MD, USA). The cells were cultured in a T-75 tissue culture flask (Falcon; Franklin Lakes, NJ, USA) with F-12K Medium (Kaighn’s Modification of Ham’s F-12 Medium, ATCC^®^ 30-2004) supplemented with 10% heat-inactivated fetal bovine serum (FBS; GIBCO&SOL; BRL Life Technologies), penicillin (100 units mL^−1^), and streptomycin (100 μg/mL). Fibroblasts were used between the 4th and 10th passage. Confluent fibroblasts were trypsinized (Trypsin-EDTA; GIBCO/BRL Life Technologies, 0.05% trypsin 0.53 mM EDTA), resuspended in serum free F-12K medium at a working concentration of 3 × 10^6^cells/mL, and kept on ice prior to use in the CGC assay.

AML14.3D10 cells (cell line was generously provided by Cassandra Paul (Wright State University, Dayton, OH)), a differentiated human myeloid leukemic cell line that displays typical morphology and enzymatic activity of normal eosinophils (Ackerman et al., 2000; Baumann and Paul, 1997; Esnault et al., 1998), were grown in T-75 tissue culture flasks in RPMI 1640 media containing 8% fetal calf serum, supplemented with 2 mM L-glutamine, 1 mM sodium pyruvate, 0.05 mg/ml gentamycin, and 5.5×10^−5^ M 2-mercaptoethanol. Eosinophil concentration was maintained between 1×10^5^ - 1×10^6^/mL in the flask. All cells were maintained in a 37 °C incubator with 5% carbon dioxide.

### HFL-1 collagen gel contraction assay in the CGC device

Two parts of HFL-1 in serum free F-12K media (3×10^6^ cells/mL) were mixed together with one part of type I collagen (9.4 mg/mL; Corning, Corning, NY, USA) and one part of 1×HEPES to a final concentration of 2.35 mg/ml of collagen, 1.5×10^6^ fibroblasts/mL. CGC devices were assembled and inserted into 24 well plates. 14 μL of fibroblast-laden collagen was loaded into the loading channel, contacting the free-swinging arm head and swinging it into the loading channel (as shown in Figure 1). 40 μg of F-12 K media supplemented with 10% FBS (for all conditions) was pipetted into the retraction tube to contact the free-swinging arm (Figure 1); the media was then quickly withdrawn to pull the free-swinging arm back into the retraction tube (Figure 1). The plate was incubated for 15 min at 37 °C. For monoculture experiments (Figure 2), 2 mL F-12 K media with or without 10% FBS was gently loaded into each well of the 24 well plate; for coculture experiments (Figure 3), 7×10^5^ eosinophils were resuspended with 2 mL of serum-free F-12K media then gently added into each well. The plate was incubated overnight.

**Figure 1:**
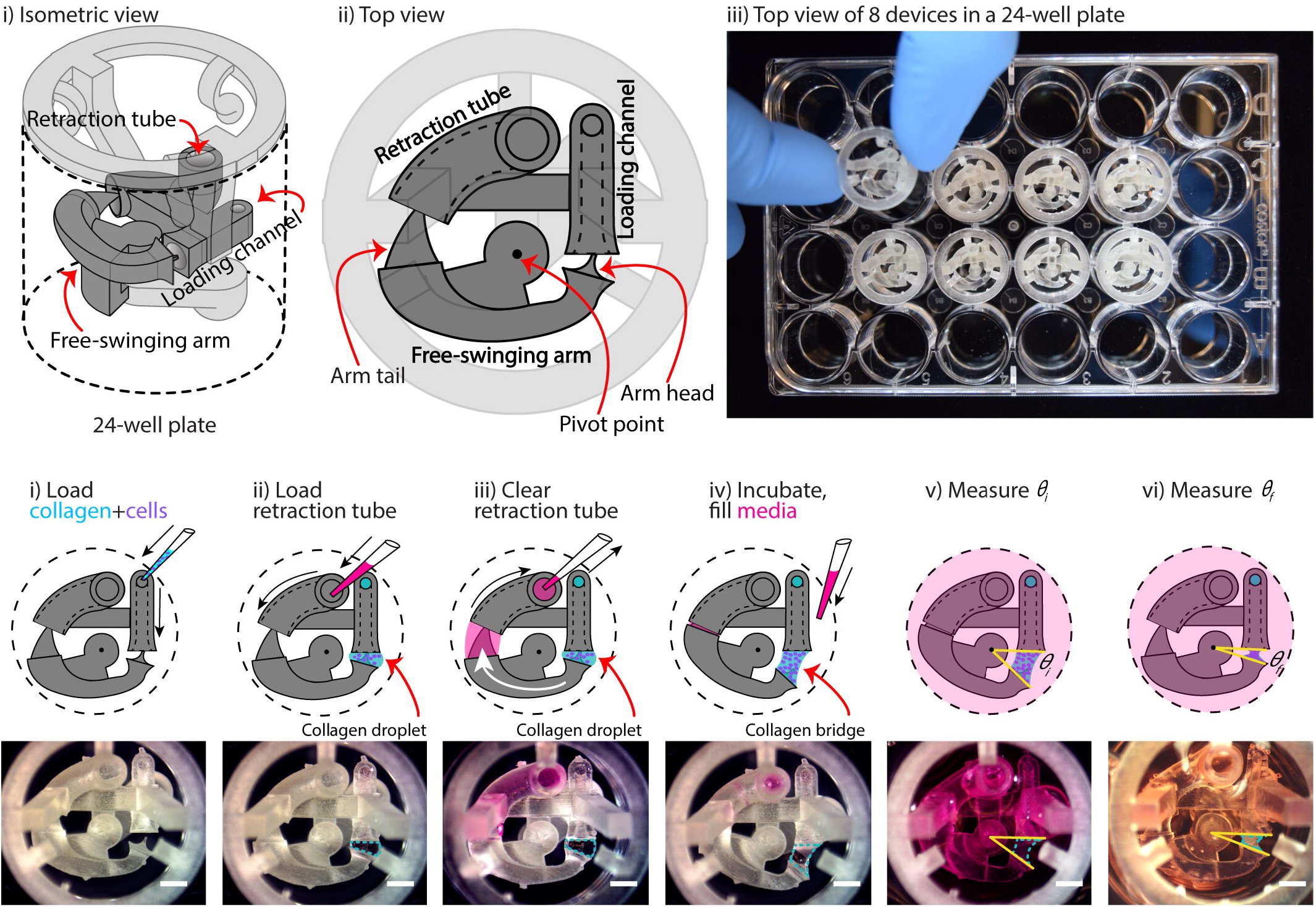
Overview of device configuration and operation. (A) Schematic drawing of an assembled collagen gel contraction (CGC) device inserted in a 24-well plate. The CGC device consists of a collagen loading channel, a free-swing arm, and a retraction tube. (B) Top view of CGC device operation work flow. i) 14 *µ*L of cell-laden collagen is loaded into the loading channel; after filling the loading channel, a collagen droplet is formed in between the loading channel and arm head. ii) and iii) 25 *µ*L of cell culture media is pipetted in and out of retraction tube; the arm tail is pulled back into the retraction tube with the flow of the media, causing the collagen droplet to extend into a collagen bridge. iv) The system is incubated at 37 °C for 15 min for collagen to gel; cell culture media is loaded directly into the well plate from top. v) and vi) The top view of the device is captured to determine the angle at starting point (*θ*_*i*_) and end point (*θ*_*f*_), respectively. The change in *θ* reflects cell contractility. Scale bars: 2 mm.

**Figure 2:**
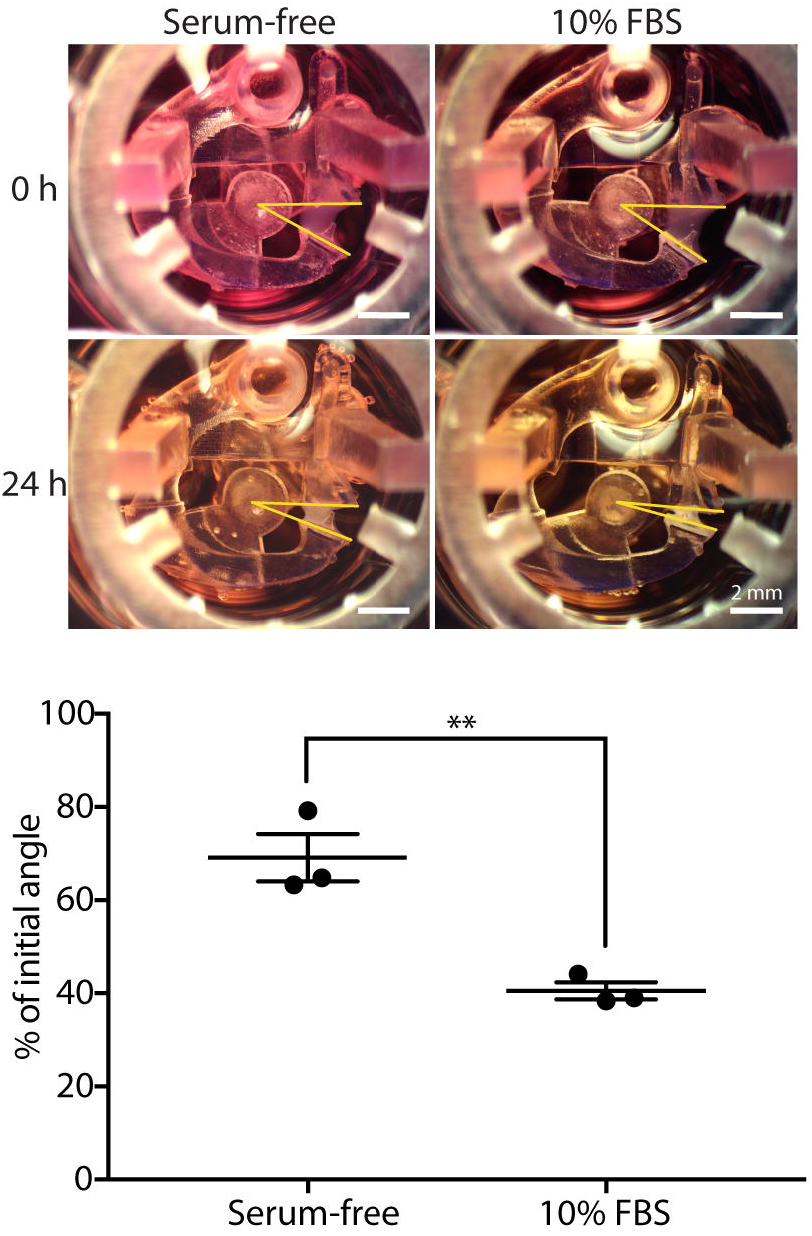
CGC device characterization using fibroblast contraction in differential serum conditions. (A) Representative images showing the contracting angle, *θ*, of the same device immediately after loading the cell-laden gel and cell culture media (0 h, top) and after 24 h in culture (bottom), in both serum-free media (left) and media containing 10% FBS (right) (scale bars: 2 mm). (b) fibroblasts (HFL-1) cultured in media containing 10% FBS contract collagen gel more than fibroblasts cultured in serum-free media. Each data point represents the average of three devices from an independent experiment; three independent experiments were performed. Error bars: SEM of three independent experiments; ** indicates significantly different values according to a two-tailed unpaired Student’s t-test (p ≤ 0.01).

**Figure 3:**
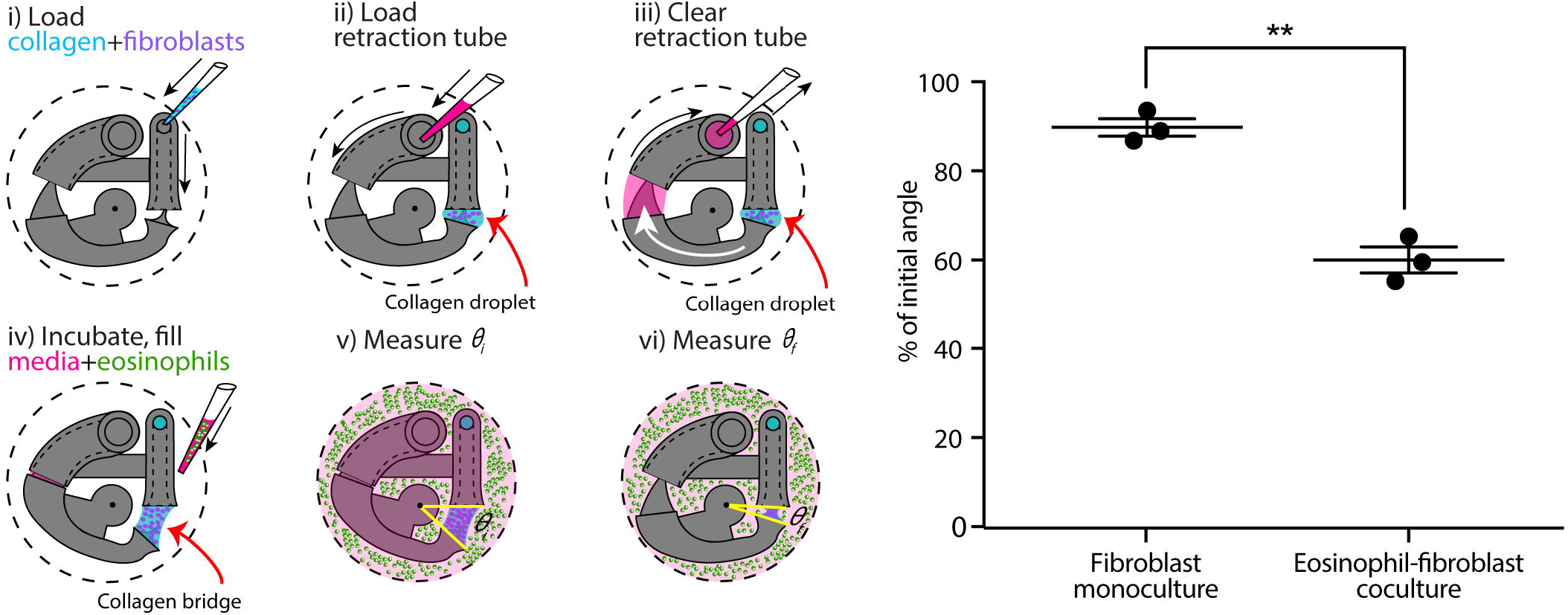
CGC device application in a coculture system with human fibroblast cells (HFL-1) and eosinophil model cell line (AML14.3D10) to evaluate the effect of soluble factor signaling from eosinophils on fibroblast gel contraction. (A) Schematics of the coculture configuration and workflow. i) Fibroblast-laden collagen is loaded into the CGC device in a 24-well plate. ii) and iii) Cell culture media is pipetted in and out of retraction tube and the arm tail is pulled back into the retraction tube. iv) The system is incubated for the collagen to gel; eosinophils are suspended in serum-free F-12K media at a concentration of 3.5 ×10^7^ cells/mL; 2 mL of serum-free media (for monoculture) or cell suspension (for coculture) is added into each well. v) and vi) Top view image is taken to measure *θ*_*i*_ and *θ*_*f*_, respectively. (B) Coculture of fibroblasts with eosinophils augments HFL-1 collagen gel contraction in serum-free media. Each data point represents the average of three devices from an independent experiment; three independent experiments were performed. Error bars: SEM of three independent experiments; ** indicates significantly different values according to a two-tailed unpaired Student’s t-test (p ≤ 0.01).

### Measurement of CGC device angle

The top view of each device was imaged using a MU1403B Microscope Camera mounted on an Amscope SM-3TZ-80S stereoscope (Amscope, Irvine, CA). Each device was imaged after setup and media addition (initial angle, *θ*_*i*_) then placed into incubator, and imaged with the same setup (position, lighting condition, and same Amscope parameters) after 24 h (final angle, *θ*_*f*_). The brightness of each image was adjusted with Fiji (ImageJ, version 2.0.0), and the CGC device angle was determined by the pivot point of the rotation axis and two side faces of collagen in contact with the device (as depicted in Figure 1b and Figure 2a). The initial and final angles were measured automatically with ImageJ for each device, respectively.

### Statistical analysis

Data are presented as the percentage of the initial angle (i.e., *θ*_*f*_/(*θ*_*i*_). Data are plotted as the mean of three independent experiments ± the standard error of the mean (SEM); each plotted point on the graphs in Figures 2 and 3 represents an independent biological experiment and is the mean of three devices within each experiment. Differences between two groups of data were evaluated using a two-tailed unpaired Student’s t-test (Prism, GraphPad Software).

## Results

### Device design and workflow

The underlying principle of our device is that it moves dynamically in response to cell-induced collagen gel contraction. As shown in Figure 1A, the device is composed of two parts: a base insert (which contains the loading channel and retraction tube) and a free-swinging arm. The free-swinging arm rests on a pivot point that juts out from the base insert, which allows the arm to rotate freely inside the well. When mechanical force is applied by cell-mediated collagen gel contraction, the device is dynamically reconfigured via rotation of the free-swinging arm, as shown in Figure 1B. The movement of the device can then be used as a quantitative metric for cell contractility by measuring the change in CGC device angle as described below. To begin the device workflow, cell-laden collagen gel precursor solution is pipetted into the loading channel of the device, where it flows through a closed tube to an opening that is positioned adjacent to the head of the free-swinging arm (Figure 1B(i)). As the gel precursor solution is added, it forms a droplet at this opening, which grows until it meets the head of the free-swinging arm. Surface tension then pulls the head of the arm into the droplet. In the next step, media is pipetted into the retraction tube of the device (Figure 1B(ii)). The media is pipetted in excess of the inner volume of the tube such that the media wets the tail of the free-swinging arm. Upon immediate withdrawal of the media in the retraction tube (which does not remove all of the media in the retraction tube), the wetted arm tail is pulled into the retraction tube by surface tension (Figure 1B(iii)). The resulting surface tension force at the arm tail pulls the spherical droplet into a hyperboloid, or bridge, at the arm head. When the device is incubated at 37 °C, the collagen bridge gels, setting the three-dimensional geometry of the bridge, and the well is filled with media (Figure 1B(iv)). The surface tension force exerted by the media in the retraction tube is nullified because the surface that applies the force is submerged in media; this allows the device to move freely in response to the force that the cells exert when contracting the gel. Finally, the initial angle (*θ*_*i*_) is measured (Figure 1B(v)), and the final angle (*θ*_*f*_) is measured after 24 hours of incubation (Figure 1B(vi)).

Quantification of gel contraction in our assay is done by comparing the initial angle, *θ*_*i*_(as shown in Figure 1B(v)), with the final angle, *θ*_*f*_(as shown in Figure 1b(vi)), after a period of incubation (typically 24 h) during which the cells contract the collagen gel. Importantly, our device design maximizes the dynamic range of our CGC assay measurement by forming the hyperboloid collagen bridge (through the use of the retraction tube as discussed in Figure 1B(ii, iii)), which increases the initial angle (*θ*_*i*_) that the device takes and increases the maximum potential change in angle that can occur due to cellular forces contracting the gel. Additionally, retraction of the collagen bridge is not affected by gel-substrate friction due to the dynamic nature of our device.

### Viability test

We evaluated the viabilities of fibroblasts and eosinophils in the coculture system after 24 h of incubation. Images from a live/dead stain are included in the ESI (Figure S1(iii)).

### FBS augments fibroblast gel contraction in our device

We used a simple fetal bovine serum (FBS) stimulation experiment as a proof-of-concept to validate that the microscale platform is capable of quantifying collagen gel contraction due to a known treatment; the comparison between fibroblast-mediated gel contraction under serum-free and serum-containing conditions is frequently used in macroscale CGC assays as a validation experiment (Lijnen et al., 2001, Zhu et al., 2001). Human fetal lung fibroblast (HFL-1)-laden collagen was loaded into the CGC device in both 10% FBS and serum-free media conditions. The initial angle of the CGC device (*θ*_*i*_, as shown in Figure 1B(v)), was measured immediately after cell culture media was added into each well. After 24 h of incubation, the angle was measured again (*θ*_*f*_, as shown in Figure 1B(vi)). The difference in the gel contraction can be clearly seen by eye (Figure 2A). In the absence and presence of FBS, the CGC device angle decreased to 69% and 41% of initial angle, respectively (Figure 2B). The gel contraction was reported as the average of three independent experiments performed on different days; the presence of FBS in the media resulted in significantly more collagen gel contraction than in serum-free conditions. Collagen without fibroblasts was loaded into the device as a negative control, and there was no significant change in the angle (Figure S2).

### Eosinophils cocultured with fibroblasts augment collagen gel contraction in our device

Following a similar CGC device loading workflow, we established an eosinophil-fibroblast coculture using our platform to test the hypothesis that soluble factors secreted from eosinophils increase fibroblast-mediated collagen gel contraction as observed in prior work by Zagai et al. (2004) using the traditional CGC assay. We used the eosinophil cell line model AML14.3D10, a differentiated human myeloid leukemic cell line that displays typical morphology and enzymatic activity of normal eosinophils (Ackerman et al., 2000, Baumann and Paul, 1997). Eosinophils were maintained in RPMI media supplemented as described in the Materials and Methods section (as recommended for this cell line) and resuspended into serum-free F-12K media prior to loading to the well plate for coculture (Figure 3A). Eosinophils were loaded into the culture media surrounding the device after the fibroblast-laden collagen was established in the device (Figure 3A (iv)); this setup allows us to study soluble-factor mediated signaling while keeping the eosinophils and fibroblasts physically separate from each other. The CGC device angle was measured at the starting point of the culture and after 24 h incubation as described previously. In the absence and presence of eosinophils, the CGC device angle decreased to 90% and 60% of initial angle, respectively (Figure 3B). The gel contraction was reported as the average of three independent experiments performed on different days; the presence of eosinophils resulted in significantly more collagen gel contraction than in monoculture. Differences were observed between the absolute values of percentage of initial angle in the monoculture serum-free conditions across Figures 2 and 3, likely due to higher passage number cells used in Figure 3; we discuss this further in the Discussion section. Collagen without fibroblasts was loaded into the device with the presence of eosinophils as a negative control for coculture, and there was no significant change in the angle (Figure S2).

## Discussion

The 3D culture of fibroblasts in native type I collagen gels has enabled researchers to integrate cell behaviors with surrounding matrix components, capturing some key aspects of cell-extracellular matrix interactions that are lost in simple 2D culture on plastic substrates (Duval et al., 2017; Bhatia and Ingber, 2014). Traditional CGC assays serve as a gold standard for cell contractility measurements, and we identified three ways in which we could build on and improve traditional CGC assays with our dynamic microscale system: 1) reduced cell and gel consumption, 2) straightforward measurements of collagen gel contraction that are not dependent on quantifying irregular gel shapes, and 3) the ability to perform coculture experiments to study how paracrine signaling (soluble-factor mediated signaling) between fibroblasts and other cell types affects fibroblast contractility. In this study, we developed a dynamic microscale CGC platform that integrates these criteria through use of a small gel volume (14 µL in comparison to over 100 µL typically used in traditional CGC assays (Gullberg et al., 1990; Timpson et al., 2011)), a simplified quantitative readout (CGC device angle), and compatibility with coculture. Importantly, existing cell-based assays developed around the traditional CGC assay can be readily translated to our microscale CGC assay because both assays share similar protocols for cell-laden collagen preparation. Prior systems for microscale gel droplet fabrication have achieved impressive advances in terms of miniaturization, which often involve engineering with new biomaterials or reagents such as polyethylene glycol or dextran that allow for rapid polymerization of droplets and careful management of evaporation (Moraes et al., 2013; Leung et al., 2015). Since the gel polymerization conditions are pH-sensitive and thermally driven, the addition of new materials in the process requires intensive testing of precise gelling conditions (Forgacs et al., 2003). Moreover, washing steps are required to remove the additional materials from the droplet, increasing the time and labor involved in the fabrication process (Moraes et al., 2013; Leung et al., 2015). Our CGC device serves as an alternative surface-tension driven method to manufacture microscale gel droplets (and ultimately stretch the droplets into hyperboloid bridges), eliminating the possible complications involved in adding new materials.

Gel area quantification has been a hurdle to the accuracy and reproducibility of the CGC assay largely due to the difficulties in characterization of gel boundaries, as well as aberrancies in gel shape post-contraction (Chen et al., 2013; Chapuis and Agache, 1992). Although atomic force microscopy (AFM) and traction force microscopy (TFM) serve as alternative quantification methods for cell contraction force measurement bypassing the gel border delineation issue, the instruments are normally unfamiliar to common users in traditional biological laboratory settings (Schierbaum et al. 2019). The CGC device we present in the study measures the change in CGC device angle, which converts the 3D change in gel volume to single parameter that is straightforward to measure. Three reference points are clearly and easily identified in digital images of the device, and the angle is calculated automatically using image processing software (see Methods section and Figures 1B and 2A).

Previous studies have identified a group of soluble factors including transforming growth factor-*β* (TGF-*β*), that contribute to fibroblast myodifferentiation, leading to increased expression of α-smooth muscle actin and a contractile phenotype typified by increased collagen gel contraction (Kendall and Feghali-Bostwick, 2014; Hinz et al., 2007; Grinnell et al., 2000). Myodifferentiation and fibrosis are particularly important in airway remodeling and asthma as they can lead to exacerbated symptoms and progressive damage (Hinz et al., 2007; Grinnell et al., 2000). To better understand how fibroblasts are affected by soluble factors from other types of cells in airway remodeling, researchers have conducted mixed coculture (embedding additional cell types into the fibroblasts-laden collagen), conditioned media coculture (feeding fibroblast-laden gel matrix with supernatants collected from other types of cells), and segregated Transwell coculture CGC experiments (Wygrecka et al., 2013; Fredriksson et al., 2003; Margulis et al., 2008; Zagai et al. 2007). These studies revealed that mast cells, red blood cells, and eosinophils can promote fibroblast contraction in mixed 3D coculture, or in conditioned media culture through paracrine signaling; whereas blood monocytes and lung epithelial cells attenuate fibroblast-mediated gel contractility, an important aspect of tissue repair (Sköld et al., 2000; Epa et al., 2015).

Given the importance of paracrine signaling in myodifferentiation illuminated by prior work, we developed an eosinophil-fibroblast paracrine signaling coculture model to demonstrate the ability to conduct coculture experiments with our microscale CGC assay (Figure 3A). Our coculture system enabled eosinophils, which settled to the bottom of the well plate physically separate from the fibroblast-laden gel suspended in our device above, to communicate with fibroblasts through shared media. In contrast to culture systems that involve transfer of conditioned media from one cell type to another, the shared media in our coculture model enables bidirectional signaling and signaling based on short-lived factors that may degrade in conditioned media studies (Ruth et al., 1999). Here, we cocultured HFL-1 with AML14.3D10, which is a well-characterized mature eosinophil surrogate (Figure 3A) (Baumann and Paul, 1998). Previous studies have identified eosinophil cationic protein as an important biomarker for airway inflammation, which is largely released from mature eosinophils (Koh et al., 2007; Zagai et al., 2004, 2007). Using our microscale CGC device, we found that the presence of eosinophils caused significantly increased fibroblast contractility (Figure 3B), which agreed with the previous mixed coculture experiments where eosinophils were mixed in with fibroblasts in collagen gel (Zagai et al., 2004).

It is worth noting that we observed a decrease in HFL-1 contractility (in the monoculture, serum-free condition) in the second set of experiments (Figure 3B, *θ*_*f*_/(*θ*_*I*_ = 90 ± 2%) compared to the first set of experiments (Figure 2B, *θ*_*f*_/(*θ*_*I*_ = 60 ± 3%), as the cells were at higher passage numbers in the second set of experiments. Thus, it is important to set up separate controls within each set of experiments (as we did in Figures 2 and 3); the comparison between the treatment and the control within experiments should be considered rather than the absolute value of the percentage of initial angle, which can vary across passage numbers.

In conclusion, this paper presents a novel platform that translates a traditional CGC assay to a microscale assay, minimizing fibroblast and gel consumption. Utilizing surface tension, the device enables generation of a suspended cell-laden gel with two standard pipetting steps. Gel contraction quantification is simplified to a single angle measurement. Moreover, we established an eosinophil-fibroblast coculture model using the CGC device and showed that the platform sustained segregated coculture and paracrine signaling to recapitulate aspects of immune-fibroblast-ECM interactions. Importantly, our platform captures bidirectional and time-sensitive paracrine signaling interactions which are sometimes lost in stepwise conditioned media studies due to decay of short-lived cytokines and other signaling molecules. Finally, our device fits within a standard well plate and cell culture incubator, increasing its translation to biology laboratories.

## Author Contributions

AT, EB, LD, SE, and NS conceptualized the study, using a microscale platform for gel contractility to investigate lung fibroblast contractility in the presence of inflammatory agents secreted by immune cells in asthmatic diseases. JD and TZ designed and tested the device. XS, AG, and SE assisted with cell culture, protocol development, and sample preparation. TZ carried out the experiments and acquired the images. TZ, JD, and AT wrote the manuscript. All authors reviewed the manuscript.

## Acknowledgments

This work was supported by NIH 1R35GM128648, P01 HL088594 and P01 HL08859, and the University of Washington. We thank Cassandra Paul for providing AML 14.3D10 cells, and Sam Berry for his suggestions on manuscript revision.

## References

Ackerman SJ, Du J, Xin F, Dekoter R, McKercher S, MakI R, et al. Eosinophilopoiesis: To be or not to be (an eosinophil)? That is the question: transcriptional mechanisms regulating eosinophil genes and development. Respiratory Medicine. 2000;94(11):1135–8.

Baumann MA, Paul CC. The AML14 and AML14.3D10 cell lines: a long-overdue model for the study of eosinophils and more. Stem Cells. 1998;16(1):16–24.

Bell E, Ivarsson B, Merrill C. Production of a tissue-like structure by contraction of collagen lattices by human fibroblasts of different proliferative potential in vitro. Proc Natl Acad Sci U S A. 1979;76(3):1274–8.

Bhatia SN, Ingber DE. Microfluidic organs-on-chips. Nat Biotechnol. 2014 Aug;32(8):760–72.

Chapuis JF, Agache P. A new technique to study the mechanical properties of collagen lattices. J Biomech. 1992;25(1):115–20.

Chen HC, Yang TH, Thoreson AR, Zhao C, Amadio PC, Sun YN, et al. Automatic and Quantitative Measurement of Collagen Gel Contraction Using Model-Guided Segmentation. Meas Sci Technol. 2013 Aug;24(8):0233/24/8/085702.

Dallon JC, Ehrlich HP. A review of fibroblast-populated collagen lattices. Wound Repair Regen. 2008;16(4):472–9.

Duval K, Grover H, Han LH, Mou Y, Pegoraro AF, Fredberg J, et al. Modeling Physiological Events in 2D vs. 3D Cell Culture. Physiology (Bethesda). 2017 Jul;32(4):266–77.

Epa AP, Thatcher TH, Pollock SJ, Wahl LA, Lyda E, Kottmann RM, et al. Normal Human Lung Epithelial Cells Inhibit Transforming Growth Factor-β Induced Myofibroblast Differentiation via Prostaglandin E2. PloS one. 2015;10(8): e0135266.

Esnault S, Jarzembowski JA, Malter JS. Stabilization of granulocyte-macrophage colony-stimulating factor RNA in a human eosinophil-like cell line requires the AUUUA motifs. Proc Assoc Am Physicians. 1998;110(6):575–84.

Forgacs G, Newman SA, Hinner B, Maier CW, Sackmann E. Assembly of collagen matrices as a phase transition revealed by structural and rheologic studies. Biophys J. 2003;84(2):1272–80.

Fredriksson K, Lundahl J, Palmberg L, Romberger DJ, Liu XD, Rennard SI, et al. Red blood cells stimulate human lung fibroblasts to secrete interleukin-8. Inflammation. 2003 Apr;27(2):71–8.

Grinnell F. Fibroblast biology in three-dimensional collagen matrices. Trends Cell Biol. 2003 May;13(5):264–9.

Grinnell F. Fibroblasts, myofibroblasts, and wound contraction. J Cell Biol. 1994 Feb;124(4):401–4.

Gullberg D, Tingstrom A, Thuresson AC, Olsson L, Terracio L, Borg TK, et al. Beta 1 integrin-mediated collagen gel contraction is stimulated by PDGF. Exp Cell Res. 1990 Feb;186(2):264–72.

Hinz B, Phan SH, Thannickal VJ, Galli A, Bochaton-Piallat ML, Gabbiani G. The myofibroblast: one function, multiple origins. Am J Pathol. 2007 Jun;170(6):1807–16.

Ilagan R, Guthrie K, Quinlan S, Rapoport HS, Jones S, Church A, et al. Linear measurement of cell contraction in a capillary collagen gel system. BioTechniques. 2010 Feb;48(2):153–5.

Jin T, Li L, Siow RCM, Liu K. A novel collagen gel-based measurement technique for quantitation of cell contraction force. J R Soc Interface. 2015;12(106).

Jonas O, Duschl C. Force propagation and force generation in cells. Cytoskeleton (Hoboken). 2010 Sep;67(9):555–63.

Kendall RT, Feghali-Bostwick C. Fibroblasts in fibrosis: novel roles and mediators. Frontiers in pharmacology. 2014; 5:123.

Koh GC, Shek LP, Goh DY, Van Bever H, Koh DS. Eosinophil cationic protein: Is it useful in asthma? A systematic review. Respir Med. 2007;101(4):696–705.

Leung BM, Moraes C, Cavnar SP, Luker KE, Luker GD, Takayama S. Microscale 3D collagen cell culture assays in conventional flat-bottom 384-well plates. J Lab Autom. 2015 Apr;20(2):138–45.

Lijnen P, Petrov V, Fagard R. In vitro assay of collagen gel contraction by cardiac fibroblasts in serum-free conditions. Methods Find Exp Clin Pharmacol. 2001 Sep;23(7):377–82.

Margulis A, Nocka KH, Wood NL, Wolf SF, Goldman SJ, Kasaian MT. MMP dependence of fibroblast contraction and collagen production induced by human mast cell activation in a three-dimensional collagen lattice. Am J Physiol Lung Cell Mol Physiol. 2009 Feb;296(2):236.

Mikami Y, Matsuzaki H, Takeshima H, Makita K, Yamauchi Y, Nagase T. Development of an In Vitro Assay to Evaluate Contractile Function of Mesenchymal Cells that Underwent Epithelial-Mesenchymal Transition. Journal of visualized experiments: JoVE. 2016(112):53974.

Moraes C, Simon AB, Putnam AJ, Takayama S. Aqueous two-phase printing of cell-containing contractile collagen microgels. Biomaterials. 2013 Dec;34(37):9623–31.

Redden RA, Doolin EJ. Collagen crosslinking and cell density have distinct effects on fibroblast-mediated contraction of collagen gels. Skin Res Technol. 2003 Aug;9(3):290–3.

Ruth JH, Esnault S, Jarzembowski JA, Malter JS. Calcium ionophore upregulation of AUUUA-specific binding protein activity is contemporaneous with granulocyte macrophage colony-stimulating factor messenger RNA stabilization in AML14.3D10 cells. Am J Respir Cell Mol Biol. 1999 Nov;21(5):621–8.

Schierbaum N, Rheinlaender J, Schäffer TE. Combined atomic force microscopy (AFM) and traction force microscopy (TFM) reveals a correlation between viscoelastic material properties and contractile prestress of living cells. Soft Matter. 2019;15(8):1721–9.

Sköld CM, Liu XD, Umino T, Zhu YK, Ertl RF, Romberger DJ, et al. Blood monocytes attenuate lung fibroblast contraction of three-dimensional collagen gels in coculture. Am J Physiol Lung Cell Mol Physiol. 2000 Oct;279(4):667.

Timpson P, McGhee EJ, Erami Z, Nobis M, Quinn JA, Edward M, et al. Organotypic collagen I assay: a malleable platform to assess cell behaviour in a 3-dimensional context. Journal of visualized experiments: JoVE. 2011(56): e3089.

Wygrecka M, Fau DB, Kosanovic DF, Petersen FF, Taborski B FAU, - von Gerlach, von Gerlach SF, et al. Mast cells and fibroblasts work in concert to aggravate pulmonary fibrosis: role of transmembrane SCF and the PAR-2/PKC-alpha/Raf-1/p44/42 signaling pathway. The American journal of pathology JID - 0370502.

Zagai U, Dadfar E, Lundahl J, Venge P, Sköld CM. Eosinophil cationic protein stimulates TGF-beta1 release by human lung fibroblasts in vitro. Inflammation. 2007 Oct;30(5):153–60.

Zagai U, Sköld CM, Trulson A, Venge P, Lundahl J. The effect of eosinophils on collagen gel contraction and implications for tissue remodelling. Clin Exp Immunol. 2004 Mar; 135(3):427–33.

Zhu YK, Liu XD, Sköld MC, Umino T, Wang H, Romberger DJ, et al. Cytokine Inhibition of Fibroblast-Induced Gel Contraction Is Mediated by PGE2 and NO Acting Through Separate Parallel Pathways. Am J Respir Cell Mol Biol. 2001;25(2):245–53.

