## Supplementary Material for "Investigating Fibroblast-Induced Collagen Gel Contraction Using a Dynamic Microscale Platform"

**Tianzi Zhang^1†^, John H. Day^1†^, Xiaojing Su^1^, Arturo G. Guadarrama^2^, Nathan K. Sandbo^2^, Stephane Esnault^2^, Loren C. Denlinger^2^, Erwin Berthier^1^, and Ashleigh B. Theberge^1,3*^**

^1^Department of Chemistry, University of Washington, Seattle, Washington, USA

^2^Department of Medicine, University of Wisconsin School of Medicine and Public Health, Madison, Wisconsin, USA

^3^Department of Urology, University of Washington School of Medicine, Seattle, Washington, USA

*** Correspondence:**
Corresponding Author


**Methods for viability test**

After overnight incubation, live/dead labeling reagent (LIVE/DEAD® Viability/Cytotoxicity Kit for mammalian cells, Invitrogen, L3224) was added into each well to a final concentration of 0.01 mM Calcein AM (labelling reagent for live cells, green) and 2 nM Ethidiumhomodimer-1 (labelling reagent for dead cells, red). The plate was incubated for 20 min. For fibroblast imaging, the device was removed from the well plate and mounted on a 3D printed imaging support (“microscope jig” in Figure S1; original design file is included in the ESI.). Eosinophils settled on the well plate bottom were imaged directly in the well plate (Figure S1(iv)). All images were taken with a Zeiss Axiovert 200 equipped with an Axiocam 503 mono camera (Carl Zeiss AG, Oberkochen, Germany).

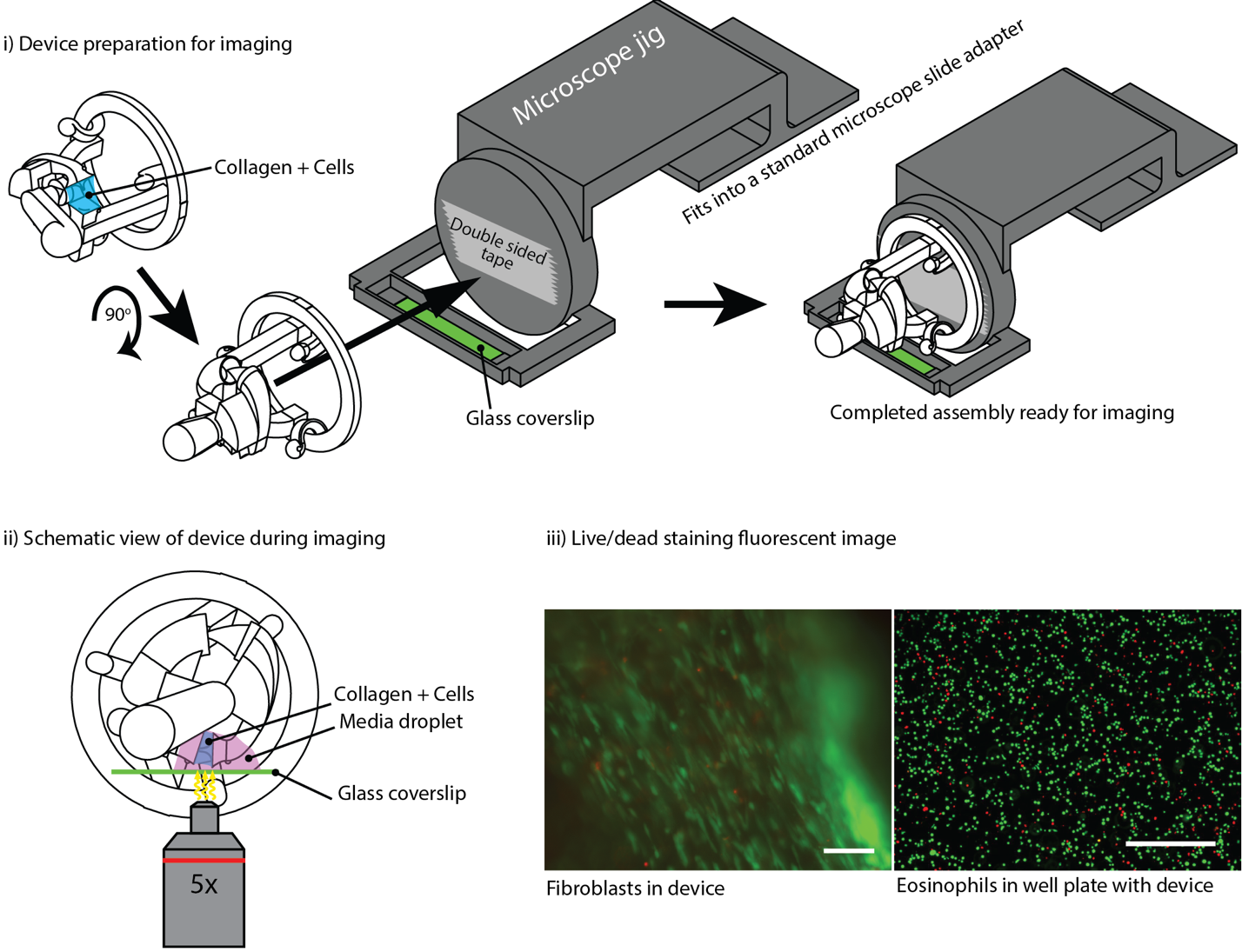

**Supplementary Figure 1**: Imaging in CGC device. (i) Imaging in device requires a microscope jig, which positions the collagen bridge directly above a glass slide. (ii) Schematic view of imaging in device. Microscope jig is omitted for simplicity. (iii) Fluorescence images of fibroblasts in collagen and eosinophils in the well plate (green: live, red: dead, scale bars: 100 μm).

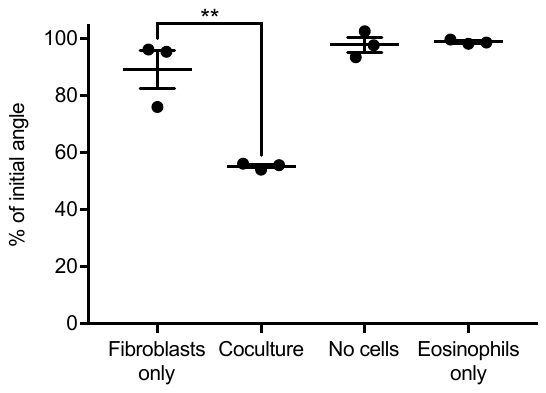

**Supplementary Figure 2**: HFL-1 mediated collagen contraction in serum-free media under four different culture conditions. “No cells” and “Eosinophils only” conditions are used as negative controls. Each data point represents a single device from the same experiment. Error bars: SEM; ** indicates significantly different values according to a two-tailed unpaired Student's t-test (p ≤ 0.01); only the “Fibroblasts only” and “Coculture” conditions were compared.
